# Dorsal raphe dopamine neurons signal motivational salience dependent on internal and external states

**DOI:** 10.1101/2020.07.27.222729

**Authors:** Jounhong Ryan Cho, Xinhong Chen, Anat Kahan, J. Elliott Robinson, Daniel A. Wagenaar, Viviana Gradinaru

## Abstract

The ability to recognize motivationally salient events and respond to them adaptively is critical for survival. Here we tested whether dopamine (DA) neurons in the dorsal raphe nucleus (DRN) contribute to this process. Population recordings of DRN^DA^ neurons during associative learning tasks showed that their activity dynamically tracks salience, developing excitation to both reward- and punishment-paired cues. The DRN^DA^ response to reward-predicting cues was diminished after satiety, suggesting modulation by internal states. DRN^DA^ activity was also greater for unexpected outcomes than for expected outcomes. Two-photon imaging of DRN^DA^ neurons demonstrated that the majority of individual neurons developed activation to reward-predicting cues but not to punishment-predicting cues, which was surprising and qualitatively distinct from the population results. Head-fixation during fear learning abolished the neural response to aversive cues, indicating modulation by behavioral context. Overall, these results suggest that DRN^DA^ neurons encode motivational salience, dependent on internal and external factors.

## Introduction

Dopamine (DA) is implicated in reward-seeking behavior and reward prediction error (RPE) encoding (Schultz et al., 1997; Bromberg-Martin et al., 2010). Increasing evidence suggests that DA also mediates non-reward functions, showing diverse responses to surprising, novel, or aversive events (Menegas et al., 2017; de Jong et al., 2019; Robinson et al., 2020; Lutas et al., 2020). These observations lead to the hypothesis that DA supports motivational control via at least two functional cell-types: one that encodes motivational value and another that signals motivational salience, defined as the absolute of motivational value (Bromberg-Martin et al., 2010). DA neurons in the lateral ventral tegmental area (VTA) or medial substantia nigra pars compacta (SNc) and those projecting to the lateral nucleus accumbens (NAc) are activated by rewarding events/cues and inhibited by aversive ones, supporting motivational value encoding (Matsumoto and Hikosaka, 2009; de Jong et al., 2019). By contrast, DA neurons in the lateral SNc and amygdala-projecting VTA cells are activated by both rewarding and aversive events/cues, consistent with salience encoding (Matsumoto and Hikosaka, 2009; Menegas et al., 2017; Lutas et al., 2020). We and others have characterized DRN^DA^ neurons, demonstrating that their population activity reflects salience rather than value (Cho et al., 2017; Groessl., 2018; Lin et al., 2020). Here, we further examine the hypothesis that DRN^DA^ neurons encode motivational salience, at both population and single-cell levels, using associative learning tasks in which the motivational salience and value of innately neutral cues were dynamically modulated by pairing them with positive, neutral, or negative outcomes (Figure 1 – figure supplement 1). We also investigated whether DRN^DA^ responses to the same motivationally salient stimuli are modulated by internal state, expectation, and/or external behavioral context.

## Results and Discussion

To explore the encoding properties of DRN^DA^ neurons, bulk fluorescence from DRN^DA^ cells expressing jGCaMP7f (Dana et al., 2019) was recorded with fiber photometry as a proxy for population neural activity (Figure 1A; Figure 1 – figure supplement 2). Mice underwent three stages of associative learning (Figure 1B). First, mice were trained in reward learning, in which one auditory conditioned stimulus (CS-A) was paired with a sucrose reward (unconditioned stimulus, US) and a second stimulus (CS-B) was paired with no reward. Subsequently, mice underwent fear training, in which the previously rewarded CS-A predicted no outcome and the previously unrewarded CS-B was paired with foot-shock. Finally, mice underwent extinction training, in which both CSs were paired with no outcome. Mice discriminated the reward-predicting CS-A from the neutral CS-B, showing increased anticipatory licks after training (Figure 1C). They also learned the contingency shifts with fear training and responded appropriately, displaying increased freezing to shock-predicting CS-B (Figure 1C).

**Figure 1:**
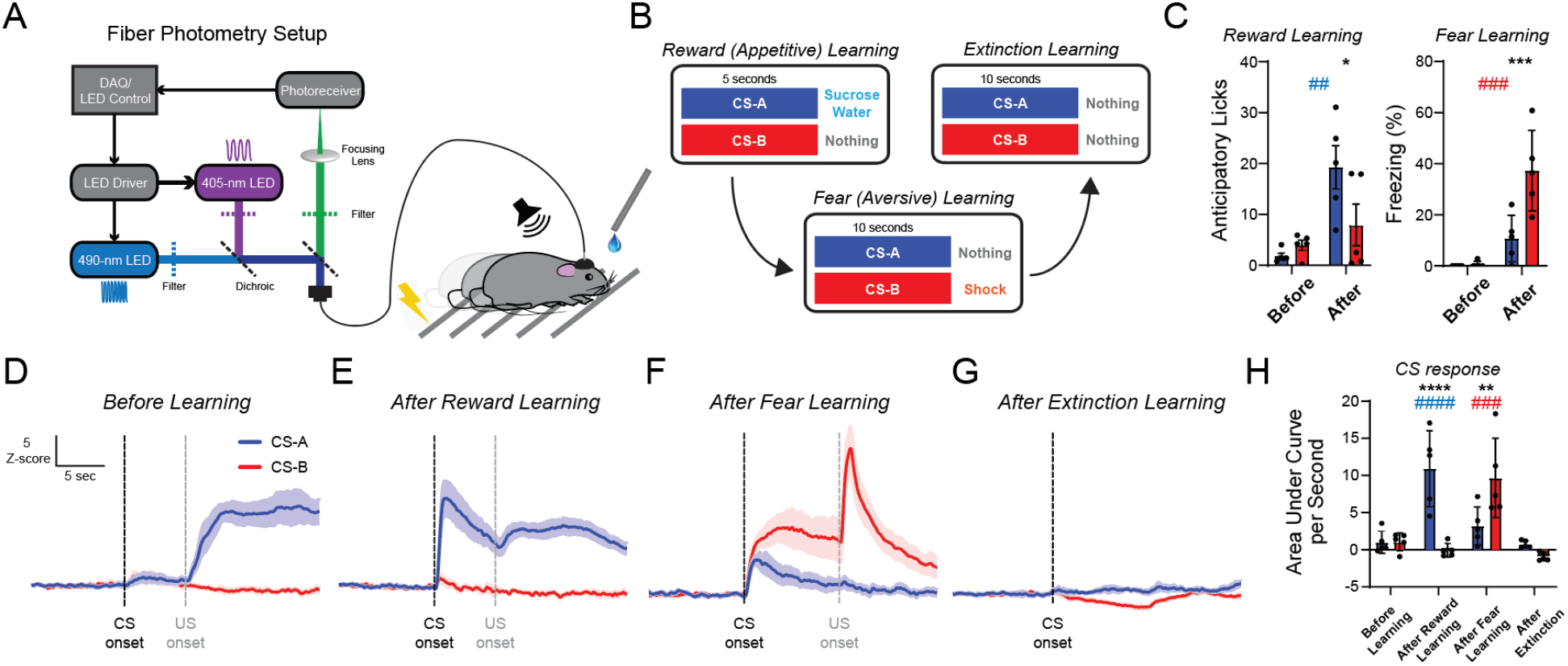
DRN^DA^ neurons dynamically track the motivational salience of conditioned stimuli. **(A)** Schematic of the fiber photometry setup used for GCaMP (490 nm) and isosbestic (405 nm) excitation and detection of emitted signals in mice freely moving in an operant chamber, which had a speaker for presenting the CS sounds, a lickometer for delivering the reward, and metal grids for delivering foot-shocks. **(B)** Three stages of associative learning with two cues (CS-A and CS-B). Reward learning was performed first, followed by fear learning and then extinction learning. **(C)** Mice successfully discriminated the CS at each stage: they showed increased anticipatory licks to CS-A (blue) after reward learning (n = 5 mice; 2-way repeated measures ANOVA; F_1,8_ = 4.583, p_time x CS_ = 0.0647; F_1,8_ = 11.54, p_time_ = 0.0094; F_1,8_ = 2.581, p_CS_ = 0.1468; *post hoc* Sidaks test; CS-A vs CS-B after learning, *p = 0.0336; before vs after for CS-A, ^##^p = 0.0089) and increased freezing behavior to CS-B (red) after fear learning (n = 5 mice; 2-way repeated measures ANOVA; F_1,8_ = 10.12, p_time x CS_ = 0.0130; F_1,8_ = 33.83, p_time_ = 0.0004; F_1,8_ = 11.09, p_CS_ = 0.0104; *post hoc* Sidaks test; CS-A vs CS-B after learning, ^***^p = 0.0006; before vs after for CS-B, ^###^p = 0.0004). **(D)** Averaged photometry response before learning for CS-A (blue) and CS-B (red), with the CS onset (black dotted line) and US onset (gray dotted line) indicated. Scale bar here also applies to (E-G). **(E)** Same as (D), but after reward learning. **(F)** Same as (D), but after fear learning. **(G)** Same as (D), but after extinction learning. Note the absence of a US onset. **(H)** DRN^DA^ neuronal response, quantified by the area under curve during cue presentation, tracks the change of salience in CS at each stage (n = 5 mice; 2-way repeated measures ANOVA; F_3,24_ = 14.98, p_time x CS_ < 0.0001; F_3,24_ = 11.89, p_time_ < 0.0001; F_1,8_ = 3.305, p_CS_ = 0.1066; *post hoc* Sidaks test; CS-A vs CS-B after reward learning, ^****^p < 0.0001; CS-A vs CS-B after fear learning, ^**^p < 0.0048; before learning vs after reward learning for CS-A, ^####^p < 0.0001; before learning vs after fear learning for CS-B, ^###^p = 0.0003). Data are presented as the mean ± S.E.M.

Photometry data showed that before learning (day 1 of reward training), CS responses were small for both CSs, followed by increased activity upon reward consumption (Figure 1D). After reward learning, the reward-predicting CS-A induced excitation whereas the response to the neutral CS-B remained small (Figure 1E). After fear learning, the CS-A response became smaller as it no longer predicted reward, and the CS-B response became larger, reflecting its pairing with the aversive US (Figure 1F). After extinction learning, both CS responses were reduced to baseline (Figure 1G). Collectively, these results suggest that DRN^DA^ population activity dynamically tracks the motivational salience of cues through increases in activity, regardless of the valence of the cue (Figure 1H and Figure 1 – supplement 1; Groessl et al., 2018; Lin et al., 2020).

The motivational salience of cues may depend on the animal’s internal state: for example, water-predictive cues are highly salient to thirsty animals but are perceived as less salient and attractive if satiated. To test this idea, after mice were fully trained in the reward-learning task, they completed 50% of trials while thirsty and the other 50% while satiated (Figure 2A). After satiety, mice stopped responding to the reward-predicting CS-A, as evidenced by the extinction of anticipatory licking (Figure 2B). Neural responses to the CS-A were also diminished after satiety (Figure 2C and 2E) while responses to the neutral CS-B remained unchanged (Figure 2D and 2E), suggesting that CS salience signals can be modulated by internal motivational states.

**Figure 2:**
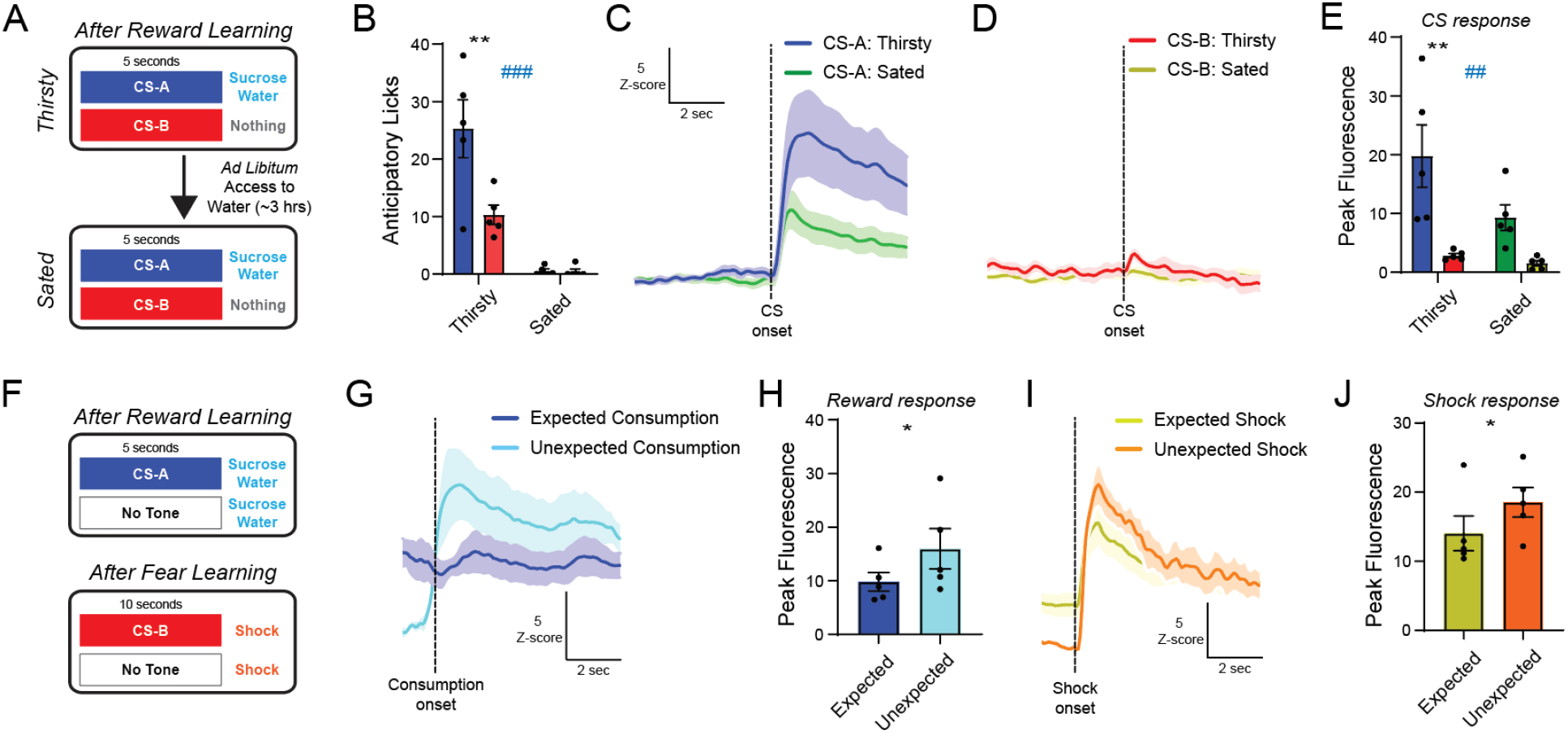
DRN^DA^ neuronal responses are modulated by internal state and expectation. **(A)** To test whether DRN^DA^ CS responses were influenced by the animals’ internal state, fully trained mice underwent half of a reward learning session while thirsty and completed the other half while sated. **(B)** Behavioral response to reward-paired CS-A, quantified by anticipatory licks during CS presentation, was reduced after satiety (n = 5 mice; 2-way repeated measures ANOVA; F_1,8_ = 8.093, p_time x CS_ = 0.0217; F_1,8_ = 43.93, p_time_ = 0.0002; F_1,8_ = 7.688, p_CS_ = 0.0242; *post hoc* Sidaks test; CS-A (blue) vs CS-B (red) when thirsty, ^**^p = 0.0022; thirsty vs sated for CS-A, ^###^p < 0.0003). **(C)** Averaged CS-A response during thirsty (blue) and sated (green) states. Scale bar here also applies to (D). **(D)** Averaged CS-B response during thirsty (red) and sated (yellow) states. **(E)** The CS-A response was significantly diminished after satiety, while the CS-B response showed no change (n = 5 mice; 2-way repeated measures ANOVA; F_1,8_ = 5.699, p_state x CS_ = 0.0440; F_1,8_ = 9.393, p_state_ = 0.0155; F_1,8_ = 11.68, p_CS_ = 0.0091; *post hoc* Sidaks test; CS-A vs CS-B during thirsty, ^**^p = 0.0015; thirsty vs sated in CS-A, ^##^p = 0.0097). **(F)** To examine whether DRN^DA^ US responses were modulated by expectation, 5% sucrose or foot-shock were occasionally introduced in the absence of predictive cues after reward and fear learning, respectively. **(G)** Averaged DRN^DA^ response to expected (dark blue) versus unexpected (light blue) reward consumption. Photometry traces were aligned to consumption onset. **(H)** Unexpected reward consumption evoked higher neural activity than expected consumption, quantified by peak fluorescence (n = 5 mice; paired t-test; t_4_ = 2.836, *p = 0.0470). **(I)** Averaged DRN^DA^ response to expected (orange) versus unexpected (yellow) shock delivery. Photometry traces were aligned to shock onset. **(J)** Unexpected foot-shock induced higher neural activity than expected shock delivery, quantified by peak fluorescence (n = 5 mice; paired t-test; t_4_ = 3.539, *p = 0.0240). Data are presented as the mean ± S.E.M.

DA neurons can be modulated by surprise or expectation, signaling prediction error (Schultz et al., 1997). To examine if DRN^DA^ neurons are modulated by prediction or expectation, mice received unexpected rewards or shocks in the absence of predictive cues, among regular CS–US pairings (separately after reward or fear training; Figure 2F). DRN^DA^ responses were larger for unexpected rewards than for expected consumption (Figure 2G and 2H). Additionally, DRN^DA^ neurons showed larger responses to unexpected shocks than expected ones (Figure 2I and 2J). Together, these suggest that DRN^DA^ neurons signal unsigned prediction errors.

DRN^DA^ neurons track the motivational salience of CSs at the population level, as demonstrated by increases in bulk fluorescence (Figure 1), but it is unclear how individual neurons are tuned to salient cues with distinct valence. Thus, we performed two-photon imaging (Figure 3A) to visualize calcium responses in single DRN^DA^ neurons. For this, gradient index lenses were implanted over the DRN to image GCaMP6m-expressing DRN^DA^ neurons (Figure 3B and Figure 3 – figure supplement 1), and mice were habituated to head-fixation for imaging. Mice underwent habituation for 35 minutes per day for 8–10 days. We imaged multiple fields-of-view (Figure 3C) while mice performed associative learning tasks (Figure 3D). Our analysis focused on the CS response, as US delivery (especially tail-shock) introduced uncorrectable motion from body movement. Before learning, only a small fraction of neurons showed a significantly increased CS response over baseline (Figure 3E). After reward learning, the majority of single DRN^DA^ cells developed increased responses to the reward-predicting CS-A only (Figure 3F). Surprisingly, after fear learning, most DRN^DA^ neurons did not show significant changes in activity from baseline, even to the shock-predicting CS-B (Figure 3G). The absence of aversive cue responses was striking, given that our and other previous results from freely-moving photometry or microendoscopic imaging showed robust increases (Figure 1F; Groessl et al., 2018; Lin et al., 2020). We reasoned that the change in behavioral context (freely-moving versus head-fixed) may have caused this unexpected variation in DRN^DA^ responses.

**Figure 3:**
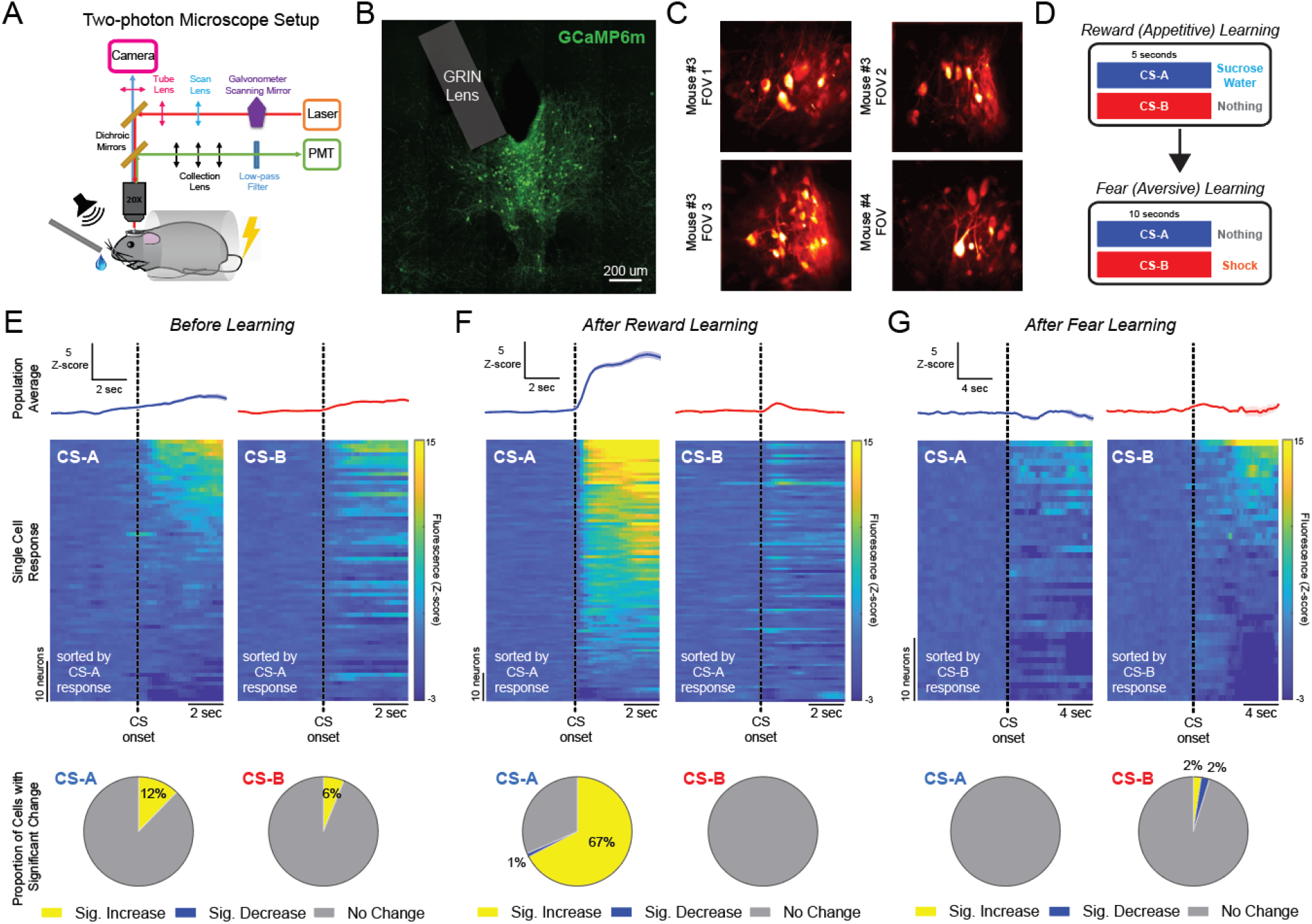
Single-cell DRN^DA^ responses to conditioned stimuli during reward and fear learning, measured by head-fixed, two-photon calcium imaging. **(A)** Schematic of the two-photon microscope setup, in which mice were head-fixed under the objective. Reward was provided through a lickometer and tail-shock was used as an aversive US. **(B)** To visualize DRN^DA^ neurons at the single cell level, AAV encoding cre-dependent GCaMP6m was injected into the DRN. Gradient index (GRIN) lenses were implanted at a 25° angle, followed by implantation of a head ring for head-fixation. **(C)** Example fields-of-view (FOVs), visualized as standard deviation projection images. **(D)** Two stages of associative learning with two cues (CS-A and CS-B). Reward learning was performed first, followed by fear learning. **(E)** DRN^DA^ neuronal responses to the CS before learning (day 1 of reward learning). Top panel: population average of all imaged cells during CS-A (blue, left) and CS-B (red, right). Middle panel: heatmap of the averaged CS responses of individual DRN^DA^ cells during CS-A (left) and CS-B (right). Neurons are sorted by the area under the curve of the CS-A response. There were 65 neurons in total, from 6 FOVs in 4 mice. Bottom panel: proportion of neurons that showed a significant increase, a significant decrease, or no change in activity in response to the CS, relative to baseline. Significance was determined by Wilcoxon sign-rank test followed by false discovery rate correction to account for multiple comparisons (q < 0.05). **(F)** Same as (E), but after reward learning. There were 95 neurons in total, from 8 FOVs in 4 mice. **(G)** Same as (E), but after fear learning. There were 42 neurons in total, from 3 FOVs in 3 mice. Data are presented as the mean ± S.E.M.

To test this hypothesis, we performed photometry in freely-moving and head-fixed mice undergoing a similar fear learning procedure (single session, 6 paired CS–US events; foot-shocks to freely-moving and tail-shocks to head-fixed mice, due to differences in experimental setups; Figure 4A and Figure 4 – figure supplement 1). All mice were water restricted and habituated for head-fixation, and then randomly assigned to one of the two groups. Freely-moving mice learned the association within these trials, showing a progressive increase of freezing in response to the CS (Figure 4B). Head-fixed mice showed a rapid decrease in licking as they received CS–US pairings (Figure 4C and 4D). CS responses in freely-moving mice gradually increased across paired trials (Figure 4E and 4G). However, head-fixed mice showed no significant change in CS responses across trials (Figure 4F and 4G). This group difference cannot be explained by the distinct shock methods, as both induced similar US responses (Figure 4 – figure supplement 2). Both groups showed learning in the form of increased freezing to the CS compared with baseline during freely-moving recall sessions, albeit with a group difference (Figure 4H). Altogether, these results indicate that salience signaling of DRN^DA^ cells can be modulated by behavioral context, especially during stressful situations where mice are forced to be immobile and receive aversive reinforcers.

**Figure 4:**
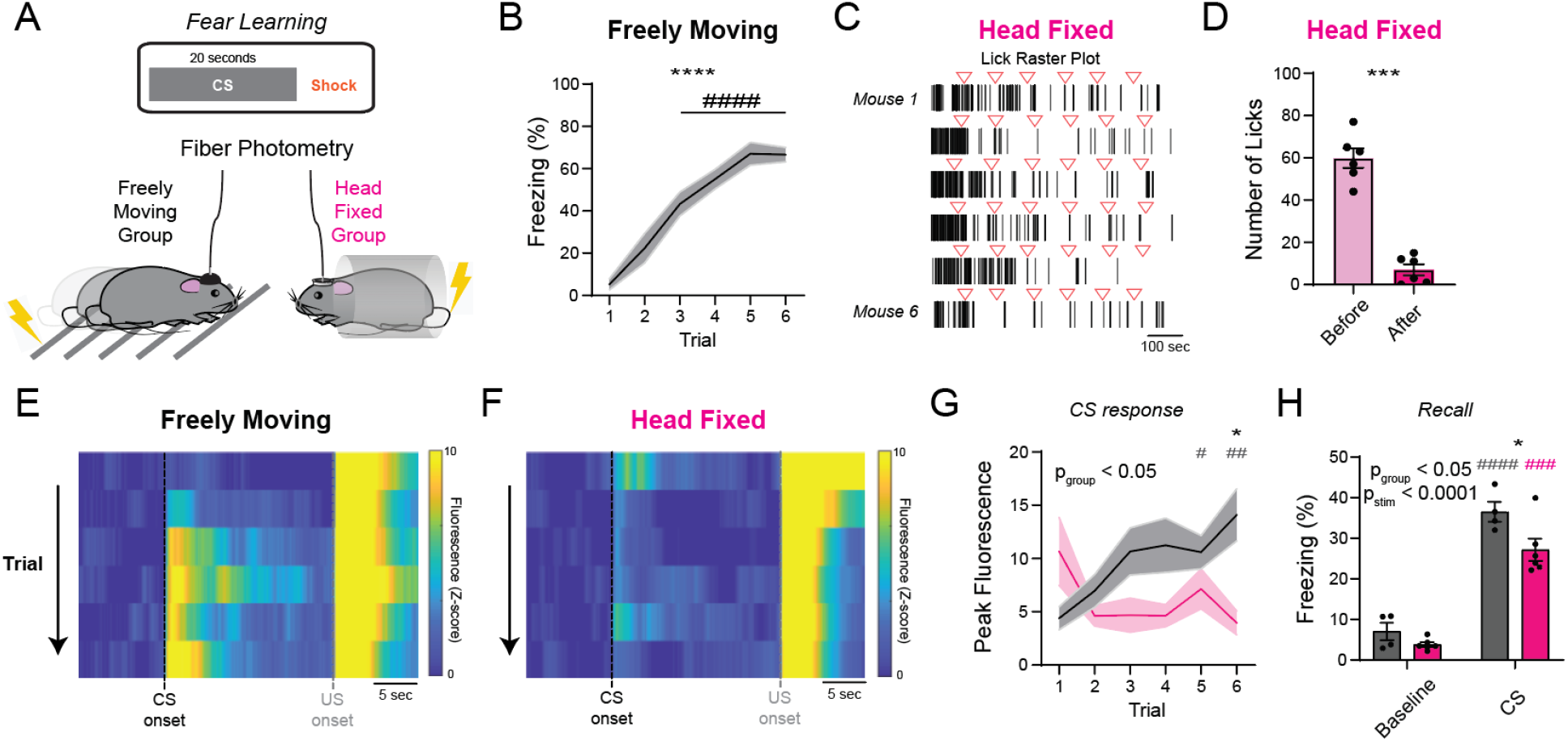
Population DRN^DA^ responses to aversive, shock-predicting CS depend on the behavioral context during fear learning. **(A)** Mice were divided into two groups (freely-moving and head-fixed) and underwent fear learning experiments. Freely-moving mice received foot-shocks as the US in an operant chamber, whereas head-fixed mice received tail-shocks. **(B)** Quantification of freezing during the shock-predicting CS showed that the mice in the freely-moving group learned the CS-US association within 6 trials (n = 10 mice; 1-way repeated measures ANOVA; F = 48.37; ^****^p < 0.0001; *post hoc* Sidak’s test; Trial 3 to 6 vs Trial 1, ^####^p < 0.0001). **(C)** Raster plot of licks from 6 head-fixed mice (each row) during fear learning. Red triangles denote the onset of shock-predictive cues. Note that, before fear learning, these mice were already habituated to the head-fixation setup with occasional sucrose delivery, so they licked continuously at the start. This licking behavior reduced dramatically across repeated CS-US pairings. **(D)** The number of licks was significantly decreased after 6 trials of fear learning compared with the baseline period prior to the first CS-US presentation (n = 6 mice; paired t-test; t_5_ = 9.817, ***p = 0.0002). **(E)** Heatmap of the averaged photometry responses in the freely-moving group across 6 trials. Each row is the average response of all mice in the group. Note the gradual development of a time-locked CS response across CS-US pairings. **(F)** Same as (E), but for the head-fixed group. Note the absence of time-locked CS response even across repeated CS-US pairings. **(G)** Freely moving mice (black) showed a significant increase in the CS response during fear learning, whereas head-fixed mice (magenta) showed no change (n = 10 freely-moving mice, n = 11 head-fixed mice; 2-way repeated measures ANOVA; F_5,95_ = 6.243, p_trial × group_ < 0.0001; F_3.302,62.73_ = 1.087, p_trial;_ = 0.3645; F_1,19_ = 4.482, p_group_ = 0.0434; *post hoc* Sidaks test; freely-moving vs head-fixed group in Trial 6, ^*^p = 0.0150; Trial 1 vs Trial 5 in freely moving group, ^#^p = 0.0452; Trial 1 vs Trial 6 in freely moving group, ^##^p = 0.0042). **(H)** Both groups showed increased freezing compared with baseline during the freely-moving recall test performed the next day (4 CS presentations in a novel arena, averaged), albeit with a group difference (n = 4 freely-moving mice; n = 6 head-fixed mice; 2-way repeated measures ANOVA; F_1,8_ = 1.639, p_stim × group_ = 0.2364; F_1,8_ = 122.3, p_stim_ < 0.0001; F_1,8_ = 10.87, p_group_ = 0.0109; *post hoc* Sidaks test; freely-moving vs head-fixed group during CS, ^*^p = 0.0149; baseline vs CS in freely-moving mice, ^####^p < 0.0001; baseline vs CS in head-fixed mice, ^###^p = 0.0001). Data are presented as the mean ± S.E.M.

The absence of neural responses to aversive cues during head-fixation (Figures 3 and 4) was striking, given that our results and previous studies have shown robust responses to aversive cues when animals are freely moving (Groessl et al., 2018; Lin et al., 2020), and given that our animals were well habituated to the setup. Head-fixation is widely used in imaging and behavioral experiments due to the need for mechanical stability or convenience, but the effects on behavior or neural activity are often assumed to be negligible. However, in rodents, head-fixation affects vocalization behavior (Weiner et al., 2016), and acute head restraint reduces the reward and cue responses of VTA DA and DRN serotonergic neurons (Zhong et al., 2017). Head-fixed mice showed higher corticosterone level over control subjects, even up to 25 days of daily training (Juczewski et al., 2020). Our findings extend these observations and demonstrate that neurophysiology can be affected by head-fixation even after habituation, especially in highly stressful and inescapable contexts.

This study builds upon previous findings that DRN^DA^ activity signals the motivational salience of cues in a learning-dependent manner, increasing in response to CSs that are paired with outcomes of either valence (Groessl et al., 2018; Lin et al., 2020), and declining with extinction. We additionally demonstrated that DRN^DA^ responses to the same CS or US can be modulated by internal state, expectation, and even external behavioral context. The dynamic nature of salience encoding by DRN^DA^ neurons may serve as a “gain control” in downstream processing in the extended amygdala (Kash et al., 2008; Groessl et al., 2018) through both fast-acting glutamate and modulatory dopamine (Matthews et al., 2016, Li et al., 2016) to orient attention towards encountered stimuli and enable the selection of appropriate behavioral responses.

## Materials and Methods

### Key resources table

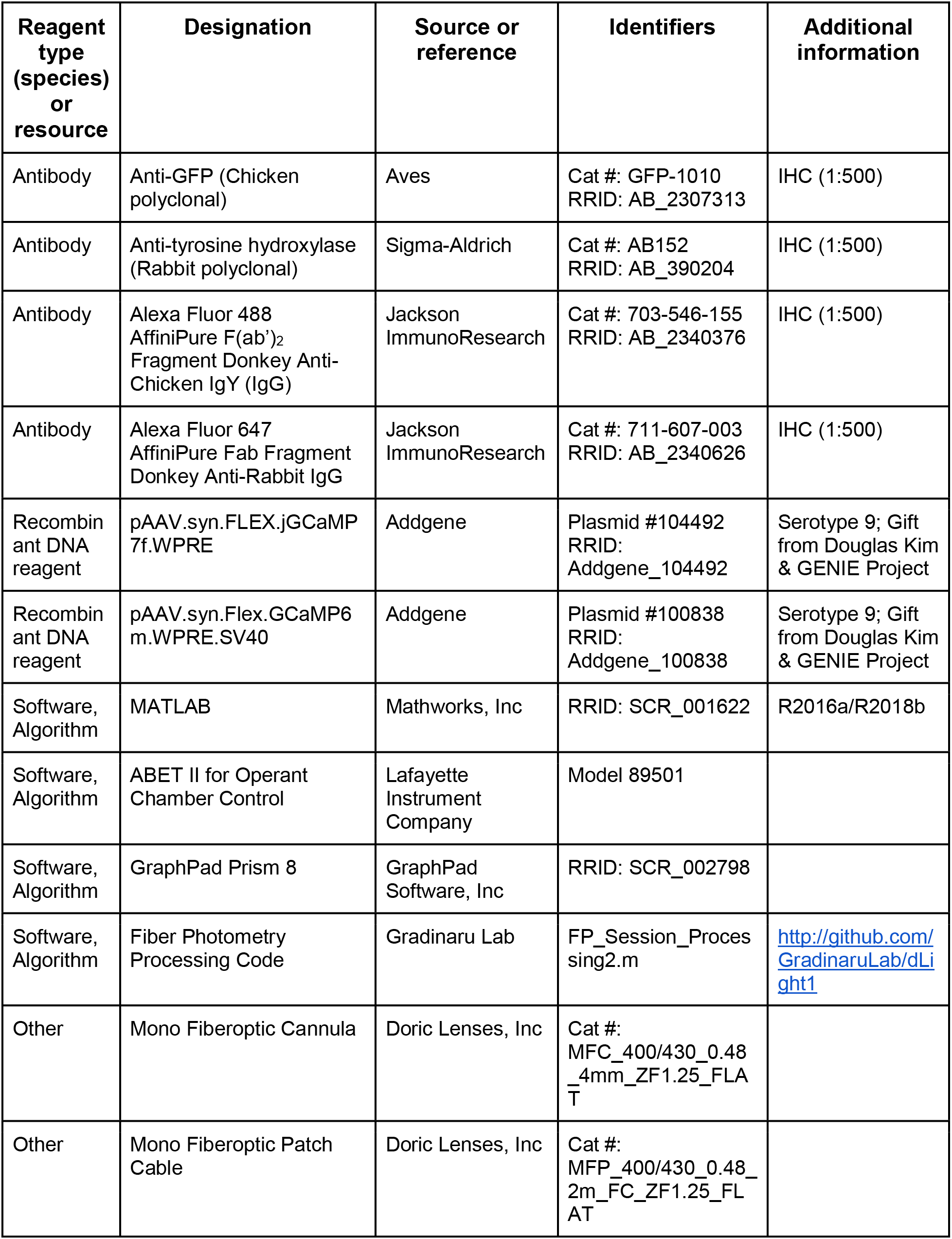
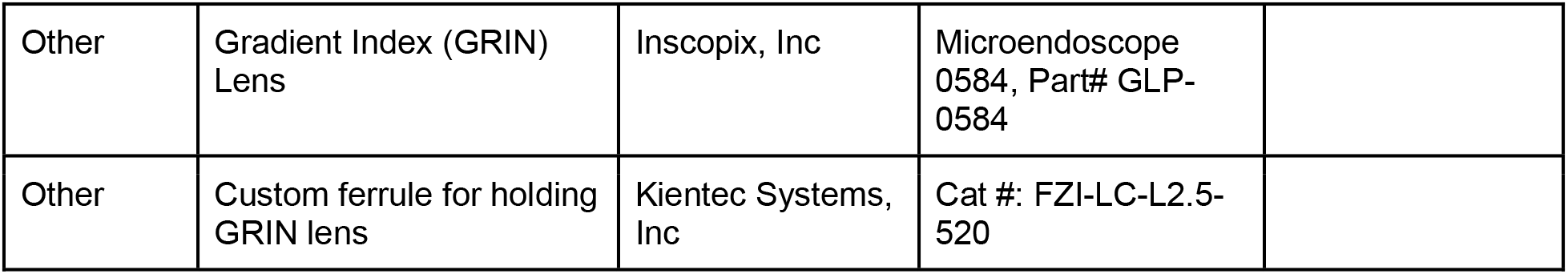

### Experimental animals

Subjects were *Th*-ires-cre transgenic mice (*Th*: tyrosine hydroxylase, a rate-limiting enzyme for dopamine synthesis; Lindeberg et al., 2004) of both sexes, aged 2–4 months at the time of surgery. *Th*-ires-cre mice were used in this study to selectively target DRN_DA_ neurons; the specificity of cre expression (compared to immunohistochemistry of *Th*+ neurons) in this mouse line has previously been shown to be around 60–75% in the DRN, which is comparable to an alternative line that express Cre recombinase under the dopamine transporter promoter (Li et al., 2016; Matthews et al., 2016; Cho et al., 2017). Typically, experiments lasted until the mice were 6–8 months old. Animals were originally group-housed but were later single-housed after undergoing surgery for photometry or two-photon imaging. Mice were housed in a room on a 12-hour light/dark cycle (lights off at 6 AM, lights on at 6 PM). All experiments were performed during the dark phase. Mice had ad libitum access to food and water before the start of water restriction (see below for details). All animal husbandry and experimental procedures involving animal subjects were conducted in compliance with the Guide for the Care and Use of Laboratory Animals of the National Institutes of Health and approved by the Office of Laboratory Animal Resources at the California Institute of Technology (IACUC protocol number: 1730). Animals were excluded from analysis if no photometry or two-photon signals were observed 4 weeks after surgery. One mouse was excluded during the associative learning experiments (Figure 1 and 2) due to health concerns related to water restriction. At the end of the experiments, the brains from all animals with fluorescence signals were histologically verified to have fibers or GRIN lenses located over the DRN.

### Surgical procedures

Stereotaxic surgeries for viral vector injections and implantation of an optical fiber/ferrule for photometry or a GRIN lens for two-photon imaging were performed as previously described (Cho et al., 2017) with slight modifications. After anesthesia (isoflurane gas/carbogen mixture, 5% for induction and 1.5–2% for maintenance), surgical preparation and exposure of the skull, a craniotomy hole was drilled in the skull (antero-posterior axis: −4.7 mm, medio-lateral axis: −1.5 mm, relative to bregma). Adeno-associated viruses (AAV) encoding jGCaMP7f or GCaMP6m in a cre-dependent manner (diluted to 1.0 × 10^13^ genome copies/mL, both from Addgene) were injected to the DRN (antero-posterior axis −4.7 mm, medio-lateral axis: −1.5, dorso-ventral axis −3.2 and −2.9 mm, relative to bregma) at a 25° angle. 300 nL of AAV was infused at each site along the dorso-ventral axis, at a rate of 50 nL per minute. After injection, the needle was held in the same place for an additional 10 minutes. Finally, the needle was slowly withdrawn over about 10 minutes to prevent backflow.

For fiber photometry, an optical fiber/ferrule (fiber: 400-μm diameter, NA 0.48, cut length: 4 mm, ferrule: 1.25-mm diameter, zirconia, glued with low-autofluorescence epoxy, Doric Lenses) was mounted to a stereotaxic cannula holder (SCH_1.25, Doric Lenses), lowered towards the DRN at a 25° angle, stopping 0.25 mm above the site of virus injection. For two-photon imaging, a 25-gauge needle (outer diameter = 0.515 mm) was attached to the stereotaxic holder (1766AP, David Kopf Instruments) and slowly lowered up to 2 mm along the dorso-ventral axis (relative to bregma, at 25°) to make a path for the GRIN lens (GLP-0584, Inscopix; 0.5-mm diameter, 8.4-mm length). Then, a small, customized zirconia ferrule (2.5-mm length, 520μm hole size; Kientec System) was carefully glued to surround the GRIN lens at one end. The same cannula holder was used to hold the GRIN lens, touching the surrounding zirconia ferrule rather than the fragile and sensitive lens. The GRIN lens was slowly lowered into the brain, stopping 0.25 mm above the site of virus injection. After implantation, a thin layer of adhesive cement was applied to the skull surface and around the implant for strong fixation. After the adhesive cement had completely dried, a layer of black dental cement was applied to build a head cap. For mice in the two-photon imaging experiments (Figure 3) and comparison of DRN^DA^ dynamics in freely-moving versus head-fixed groups (Figure 4), a customized ring for head fixation (stainless steel, 5-mm inner diameter, 11-mm outer diameter) was super-glued to the cement surface before the dental cement was fully dried, so that the ferrule or GRIN lens tip was located within the ring. More dental cement was applied inside the ring. To protect the GRIN lens from damage, the lens tip was covered with a small piece of parafilm and low-toxicity silicone adhesive (Kwik-sil, World Precision Instruments) was applied. After the silicone adhesive fully solidified, mice were unmounted from the stereotaxic frame and their recovery was monitored for about 2 hours.

### Fiber photometry

Fiber photometry was performed as previously described (Lerner et al., 2015; Cho et al., 2017; Robinson et al., 2020).

### Two-photon imaging

In vivo two-photon imaging was performed with a custom-built microscope. Briefly, a pulsed femtosecond laser beam from a Ti:Sapphire laser system (940 nm), coupled with OPA (Insight DS+, Spectra-Physics, CA), passed through a beam expander (75:50) and an iris (SM1D12C, Thorlabs), set to 3 mm. An XY galvanometer (6215H, Cambridge Technology) was placed before a pair of scan lenses (LSM54-1050, Thorlabs) and a tube lens (ITL200, Thorlabs). An 805 nm short-pass dichroic mirror (DMSP805SPL, Thorlabs) was used to allow simultaneous near-infrared (IR) visualization along with two-photon excitation. Near-IR visualization for sample localization was achieved by a 75-mm tube lens (AC508-075, Thorlabs), directed to an HDMI-output camera (HD205-WU, AmScope). A 500–700 nm reflecting dichroic mirror (T600/200dcrb, Chroma) was used to split the two-photon excitation and emission paths. A 20X/0.5 NA air objective (Olympus, UPLFLN20XP) was used, and the laser power was set to 60–80 mW. Emitted photons were passed through the collective optics (AC508-100-A, *f* = 100 mm, at z = 100 mm from BA, most convex side facing the sample; and a pair of LA1131, *f* = 50 mm at z = 150 mm and z = 156 mm from the back BA, convex sides facing each other) and a 680-nm low-pass filter (et680sp-2p8, Chroma) into the photomultiplier tube (Hamamatsu R3896). Laser intensity was controlled by the rotation of a half-lambda waveplate (Thorlabs AHWP05M-980) relative to a Glan polarizer (Thorlabs GL10-B) using a motorized rotation stage (Thorlabs PRM1/Z8). Stage XY adjustment and microscope focus was controlled by motorized linear actuators (Z825B, Thorlabs). Imaging data were acquired using an FPGA DAQ board (National Instruments 7855R) and custom-written software in Labview. An electromechanical shutter (Uniblitz VS25, Vincent Associates) was used to ensure laser safety during imaging. The imaging frame size was 194 x 194 pixels with a 4-Hz frame rate. In 3 out of 4 mice, 2 or 3 fields-of-view (FOVs) were obtained (at least 100 μm apart in the Z-direction) that showed different sets of neurons. We did not try to match or keep the same FOVs across different recording days. During each imaging session, after finding an FOV, two-photon scanning was triggered for each trial 15 seconds before the CS delivery and terminated 20 seconds after the CS delivery.

### Water restriction and habituation procedures for head-fixed experiments

All animals reported here underwent water restriction procedures (1.5 mL per day, provided at 4 pm everyday), starting from 2–3 weeks after surgery when mice had fully recovered. The water restriction was mainly to motivate the animals to engage in reward learning, but the fear-learning-only cohort (Figure 4) were also water restricted to maintain consistent experimental conditions and to facilitate habituation training (see below). Once water restriction started, mice were weighed daily and were returned to ad libitum access to water if their weight loss was >10% of their pre-restriction weight. Animals were water restricted at least for 5 days before they started freely-moving associative learning tasks or habituation training for head-fixed experiments.

Regarding habituation procedures for head-fixation, we generally followed a previously published protocol (Guo et al., 2014), but some extra steps were introduced to ensure that the mice were slowly acclimated to the setup. On day 1, mice were familiarized with experimenter handling for about 15 minutes. After mice became calm on the experimenter’s hand (exhibiting grooming behavior and spending less time looking outside the hand), they were given access to up to 0.4 mL of 5% sucrose water, delivered via a 20 uL pipette. After reward consumption, mice were further handled for about 2 minutes before being returned to their home cages. On day 2, mice were handled in a similar fashion for 5 minutes with access to up to 0.1 mL of 5% sucrose water. We then introduced the body tube (made from plexiglass) with the other hand and let animals explore the tube freely. We performed this step up to 10 times until mice voluntarily entered the body tube. After entering the tube, the experimenter gently held the tail to prevent escape and mice were rewarded with up to 0.1 mL of 5% sucrose for another 5 minutes. Next, the experimenter quickly took hold of the implanted head ring and secured it to the head fixation bar (all < 10 seconds). The body tube was also secured with a lever. Mice remained head-fixed for 5 minutes and 20 uL of 5% sucrose was provided every 20 seconds. On day 3, mice were further acclimated to the apparatus, now with 10 minutes of head-fixation and 5% sucrose reward delivered every 30 seconds. From day 4 to day 7, mice were introduced to the lickometer used in the experiments and the duration of head-fixation was gradually increased from 15 to 30 minutes (increasing by 5 minutes every day), with reward provided every minute. We reasoned that mice showed good habituation and were ready to advance to the behavioral experiments when they consumed the reward throughout the duration of head-fixation and when they produced much less feces than on the first day of training. We trained mice in the head-fixation apparatus for up to 30 minutes, well above the duration of recordings (~25 minutes for two-photon imaging, ~15 minutes for head-fixed fear learning). Note that we extended the number of days of habituation training compared with Guo et al., 2014 to make sure that the mice were habituated to the experimental setting slowly, to minimize the level of stress as much as possible.

### Associative learning tasks in freely-moving photometry recordings

All behavioral experiments were programmed and executed with ABET II software (Lafayette Neuroscience). After animals were water restricted for 5 days, they were introduced to an operant chamber (Lafayette Neuroscience) and allowed to freely explore for 30 minutes with a patch cable attached. 5% sucrose was delivered to the lick port at intervals randomly drawn from a uniform distribution of 45–75 seconds, so that mice could learn the location. Licks were counted when the infrared beam at the lick port was broken. This habituation was repeated for 2 days.

After habituation, the reward (appetitive) learning phase started. Mice were introduced to two types of conditioned stimuli (CS-A and CS-B; 5 kHz tone or white noise, 75 dB, 5 seconds, counterbalanced across animals). 25 uL of 5% sucrose reward (as the unconditioned stimulus; US) was delivered only after CS-A presentation; there was no outcome after the CS-B presentation. Within a session, 20 CS-A and reward pair trials and 10 CS-B and no outcome pair trials were given. The inter-trial interval was drawn at random from a uniform distribution of 45–75 seconds. There was a total of 21 reward learning sessions for all animals, and photometry signals were recorded on day 1 (“before learning”) and day 21 (“after reward learning”). On day 1, video was also recorded.

To examine whether DRN^DA^ cue responses are influenced by internal state, mice performed half of the trials in a reward learning session (10 CS-A and reward pairs, 5 CS-B and no outcome pairs) while they were thirsty and completed the other half after satiety. In between these two separate sessions, they were given free access to water for 3 hours in their home cages. After this experiment, mice underwent regular water restriction for two days. To study whether DRN^DA^ neurons encode positive prediction error, mice performed another experimental session in which the US was presented without the predictive CS-A. In this session, there were 10 “expected” trials (CS-A paired with the US) and 5 “unexpected” trials (US only).

Subsequently, mice underwent a fear (aversive) learning phase, in which the previously reward-predicting CS-A no longer predicted any outcome and the previously neutral CS-B was paired with an aversive foot-shock (0.5 mA for 1 second). The duration of both CSs was increased to 10 seconds so that freezing behavior could be quantified. 20 CS-B and shock pairs were presented and 10 CS-A and no outcome pairs were presented with the inter-trial intervals as described above. Photometry and video recordings (to measure freezing behavior) were performed on day 2 of fear learning (“after fear learning”). On day 3, after mice were fully trained on the fear learning task, we performed a similar prediction error experiment, but now with aversive cue CS-B and foot-shock. In this experiment, 3 unpredicted shocks were presented intermixed with 10 normal CS-B and shock pairings.

Finally, animals underwent an extinction learning phase in which both CS-A and CS-B were presented (10-second duration, 15 times each) but paired with no outcomes. This was repeated for 4 days and recording was performed on day 5 (“after extinction”).

This series of experiments was performed in two cohorts (one mouse excluded due to health concerns). Results were replicated between those two cohorts.

### Associative learning tasks in head-fixed two-photon imaging

Procedures for the associative learning tasks for two-photon imaging under head-fixation were similar to the procedures used in the freely-moving condition, with some small differences. ABET II software was also used to execute the associative learning tasks. Before the imaging experiments started, mice underwent habituation training (see above) for 8–10 days and were then transferred to the microscope imaging setup. Mice were further acclimated to the imaging setup for 2 days, receiving free reward (5% sucrose) every 90 seconds for 35 minutes. “Before learning” recordings were performed on day 1 of the reward learning phase: two mice with multiple FOVs performed two separate sessions in a single day, separated by a 6-hour interval. “After reward learning” recordings were obtained on days 18–20 after the mice showed clear discrimination between the reward-predicting CS-A and the neutral CS-B on the basis of their anticipatory licking behavior. One FOV was imaged per day. During training, 20 CS-A trials and 10 CS-B trials (5 kHz tone or white noise, 75 dB, counterbalanced) were presented with inter-trial intervals of 45–75 seconds. On imaging days, 10 CS-A trials and 10 CS-B trials were presented per session.

Fear learning was conducted similarly to the methods stated above, except that tail-shock was used as the US and imaging was performed on day 2. Tail-shock (0.5 mA for 1 second) was administered via two pre-gelled electrodes wrapped around the tail and connected to a stimulus isolator (Isostim A320R, World Precision Instruments), following Kim et al., 2016. Tail-shock was triggered by external transistor-transistor logic pulses generated by ABET II software. Due to the highly aversive nature of the US, we selected only one FOV per mouse and performed recordings once for “after fear learning”. During training, 10 CS-A trials and 20 CS-B trials were presented with inter-trial intervals of 45–75 seconds. On the imaging day, 10 CS-A trials and 10 CS-B trials were presented.

This experiment was performed in two cohorts (3 mice excluded before experiments due to absence of fluorescence signals). All individual mice showed qualitatively similar and replicable results.

### Fear learning tasks in freely-moving and head-fixed photometry recordings

To examine whether DRN^DA^ responses to aversive cues are affected by the external behavioral context (freely-moving versus head-fixed conditions), we performed photometry recordings in two different behavioral contexts. All mice underwent identical surgery, water restriction, and head-fixation habituation procedures before being randomly assigned to either the freely-moving or the head-fixed group. The fear learning task was slightly different from the ones described above and was adapted from previous studies (Groessl et al., 2018; Cai et al., 2020). White noise (20 seconds) was used as the CS. The duration of the CS was set to 20 seconds to better quantify freezing behavior as an index of learning. For the US, the freely moving group received foot-shocks within an operant chamber and the head-fixed group received tail-shocks. 6 CS–US pairs were presented with inter-trial intervals of 60–120 seconds. The next day, a subset of mice from both groups performed a fear recall experiment. Mice were introduced to a novel cylindrical cage and allowed to freely explore for about 5 minutes to habituate to the novel context. After the habituation period, 4 CSs were presented with no US to see if cue-induced freezing behavior was evoked.

This experiment was performed in two separate cohorts (n = 10 and 11 mice each). Qualitatively similar and replicable results were obtained from both cohorts and across individual mice.

### Data analysis

#### Behavior

For reward learning, we counted the number of anticipatory licks, defined as licks during the CS presentation before the reward is delivered, as a proxy for learning. For fear learning in freely-moving conditions, freezing behavior was used as an index of associative learning and was quantified visually by an observer blind to the experimental condition. For fear learning in head-fixed conditions (Figure 4), the number of licks was counted throughout the session. As these mice received 5% sucrose during the habituation procedure, all tested mice showed continuous licking as soon as they were head-fixed (Figure 4C). We measured whether this licking behavior was affected by repeated CS–US pairings (Lovett-Barron et al., 2014). We note that this does not directly reveal whether the animals have learned the association between the CS and the aversive US *per se.* Therefore, on the next day, mice from both freely-moving and head-fixed groups performed a fear recall session. We compared their freezing behavior during the baseline (after 5 min habituation but before presentation of the first CS) with that during CS presentation to test whether mice could recall the shock-paired cues.

#### Fiber photometry

Acquired photometry data files were processed with custom-written MATLAB code, as in previous studies (Lerner et al., 2015; Cho et al., 2017; Robinson et al., 2020). Signals from 490- and 405-nm excitation wavelengths were low-pass filtered at 2 Hz with zero-phase distortion. To calculate ΔF/F, a least-squares linear fit was applied to the isosbestic signal and aligned to the GCaMP signal. The fitted signal was subtracted from the 490-nm signal and subsequently divided by the fitted 405-nm signal. Fluorescence signals were then converted to robust Z-scores in each trial using the median and median absolute deviation (MAD) of the baseline, defined as a 15-second epoch before the CS presentation. Neural activity was quantified either by the area under the curve (AUC) per second (Figure 1) or the peak fluorescence (Figures 2 and 4). AUC per second was used in Figure 1 because there was a possibility of inhibited or reduced activity (Figure 1 – figure supplement 1; reflected as a negative AUC value) for paired cues, and indeed our data show small but negative values for CS-B after extinction learning (Figure 1H). In other cases, neural activity was compared using the peak fluorescence, since all showed increased fluorescence from baseline upon CS or US presentation.

#### Two-photon imaging

First, the separate imaging files (one file for each trial) were concatenated in MATLAB and saved as a tiff file. The combined movie was motion-corrected using a non-rigid registration algorithm in Suite2p (Pachitariu et al., 2017) and saved as another tiff file. This motion-corrected imaging file was loaded in ImageJ and the regions of interest (ROIs, corresponding to a single neuron) were manually drawn on the basis of the mean and standard deviation projection images (McHenry et al., 2017). Fluorescence time-series were extracted for each ROI by averaging all pixels within the ROI for each frame. To remove potential contamination from neuropil or nearby dendrites/axons, we extracted the fluorescence from a ring-shaped region (after enlarging each ROI 1.5 times and excluding the original ROI) and removing pixels in other ROIs, if any. This neuropil fluorescence was subtracted from the ROI fluorescence after being scaled by a correction factor (*cf*). Usually the correction factor is estimated as the ratio of fluorescence intensity between a blood vessel and neighboring non-ROI neuropil region; however, since we were not able to identify any blood vessels in our imaging datasets, we adopted a correction factor of 0.6, which is within the range used in previous studies (Chen et al., 2013; Cox et al., 2016). Therefore, the neuropil-corrected fluorescence (F_correct_) was calculated as: F_correct_ = F_ROI_ − *cf* × F_neuropil_, after F_ROI_ and F_neuropil_ were smoothed with a median filter (length: 4). We further normalized the fluorescence in each trial by calculating the robust Z-score using the median and MAD of the baseline, defined as a 10-second epoch before CS presentation. Neural activity was quantified for each cell by calculating the AUC per second during the baseline and the CS presentation.

### Statistical analysis

Sample sizes were determined to be comparable to previous similar studies with calcium imaging (Groessl., 2018; Lin et al., 2020). Statistical analysis was performed with GraphPad Prism 8 (GraphPad Software, Inc) or MATLAB (MathWorks). All statistical tests performed and results are stated in the figure legends and provided in detail in Source Data 1. Statistical tests were chosen according to the nature of the experiments and datasets. Paired or unpaired t-tests were performed for single-value comparisons. When analysis of variance (ANOVA; one-way or two-way repeated-measures) was performed for multiple trials or groups, *post hoc* Sidak’s test was used to correct for multiple comparisons. To examine if the DRN^DA^ cue response was significantly different from baseline at the single-cell level (Figure 3), the Wilcoxon sign-rank test was used to calculate the p-value for each cell. Then, the false-discovery rate (FDR) correction was applied (q < 0.05) to correct for multiple comparisons. When the response of a neuron was statistically significant after the FDR correction, the mean value of the cue response was compared to baseline and declared “significantly increased” if it was larger or “significantly decreased” if it was smaller. No outliers were removed from any of statistical analyses.

### Histology

Mice were euthanized with CO_2_ and transcardially perfused with 15 mL of ice-cold 1x PBS with heparin (10 U/mL), and then 30 mL of ice-cold 4% PFA. Mouse brains were removed from the skull and post-fixed in 4% PFA at 4°C overnight. The PFA solution was switched to 1x PBS the next morning. Brains were cut into 50 μm coronal sections with a vibratome (VT1200, Leica Biosystems). Sections were stored in 1x PBS solution at 4°C until further processing. For immunohistochemistry, brain sections were first incubated in a 1x PBS solution with 0.1% Triton-X and 10% normal donkey serum (NDS) with primary antibodies at 4°C overnight. The next day, sections were thoroughly washed four times in 1x PBS (15 minutes each). Then brain sections were transferred to 1x PBS with 0.1% Triton-X and 10% NDS with secondary antibodies and left overnight at 4°C. The next morning, sections were washed as described above and mounted on glass microscope slides (Adhesion Superfrost Plus Glass Slides, Brain Research Laboratories). After the sections were completely dry, they were cover-slipped after applying DAPI-containing mounting media (Fluoromount G with DAPI, eBioscience). Fluorescent images were obtained with either a confocal (LSM 880, Carl Zeiss) or a fluorescence microscope (BZ-X, Keyence).

### Data and materials availability

Codes for fiber photometry signal extraction and processing are available at http://github.com/GradinaruLab/dLight1. Source data with statistical results is available as a separate excel file.

## Acknowledgment

We appreciate the entire Gradinaru lab for critical feedback. This work is supported by NIH Director’s New Innovator IDP20D091182-01, PECASE, NIH/NIA 1R01AG047664-01, NIH BRAIN 1U01NS090577, Heritage Medical Research Institute, Chen Institute (to V.G.), Beckman Institute (to V.G. and D.A.W.), TCCI Chen Graduate Fellowship (to X.C.), Colvin divisional fellowship at Caltech (to A.K.) and Children’s Tumor Foundation Young Investigator Award 2016-01-00 (to J.E.R).

---

**Figure 1– figure supplement 1.**
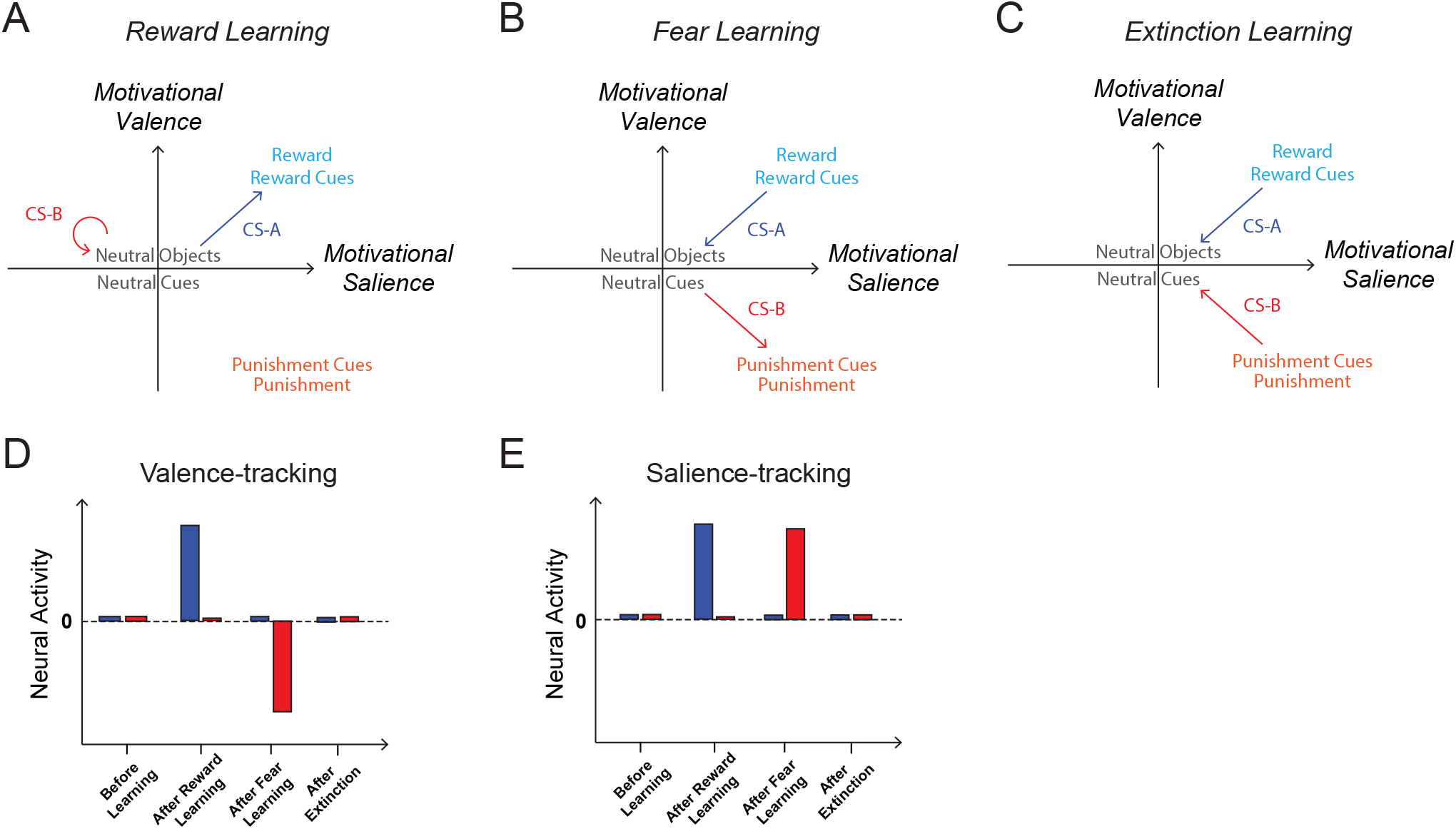
**(A)** In reward learning, the motivational valence and salience of conditioned stimulus A (CS-A) becomes positive as it is paired with reward (appetitive nature). Since CS-B is not paired with an unconditioned stimuli (US), its valence and salience stay close to zero. **(B)** In fear learning, the motivational salience and valence of CS-A, which was previously paired with reward and now has no outcome, decreases back to zero. On the other hand, as CS-B now predicts punishment (an aversive US), its valence becomes negative and its salience increases. **(C)** In extinction learning, the motivational valence and salience of both CS-A and CS-B return to zero as they no longer predict any US. **(D)** In theory, neurons that track motivational valence, such as DA neurons in the lateral VTA or those projecting to the nucleus accumbens lateral shell (Matsumoto and Hikosaka, 2009; Martin-Bromberg et al., 2010; de Jong et al., 2019), should show increased activity to reward-paired cues after reward learning and decreased activity to shock-paired cues after fear learning, compared with baseline or before learning. These changes in activity should both be reduced to close to baseline after extinction learning. **(E)** Neurons that track motivational salience should show increased activity to both reward-paired and shock-paired cues, after reward and fear learning respectively, and return to close to baseline after extinction learning.

**Figure 1 – figure supplement 2.**
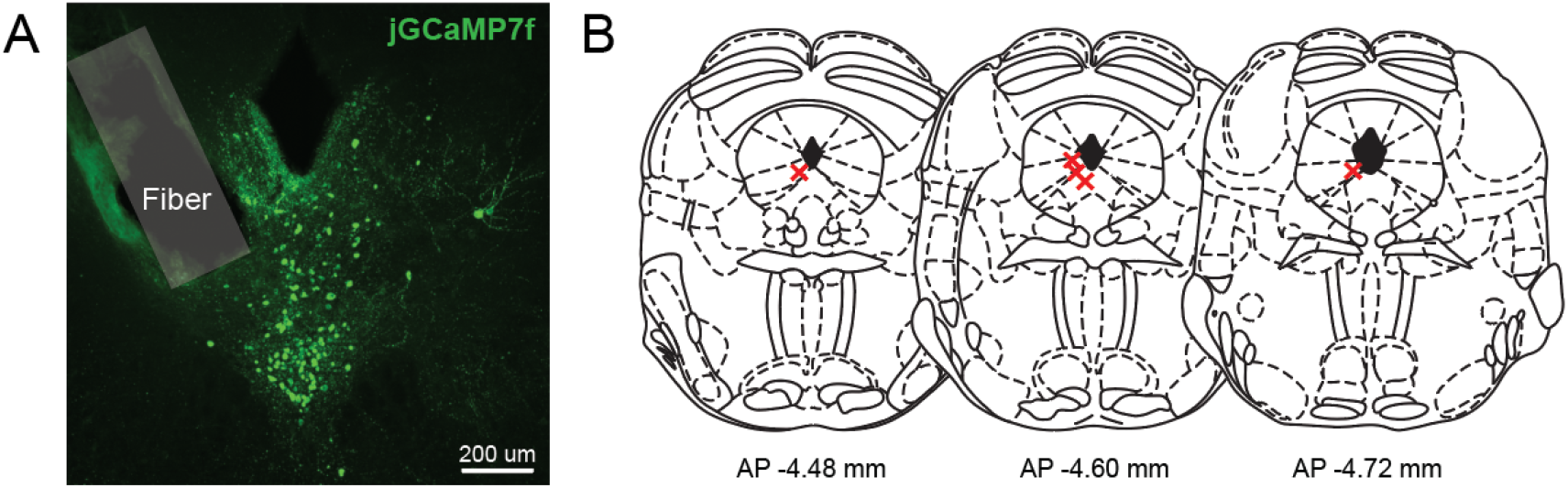
**(A)** A representative histological image of jGCaMP7f-expressing DRN^DA^ neurons showing the location of the photometry fiber tip. **(B)** Schematic of the anatomical locations of individual fiber implants in the 5 mice used in the experiments shown in Figures 1 and 2.

**Figure 3 – figure supplement 1.**
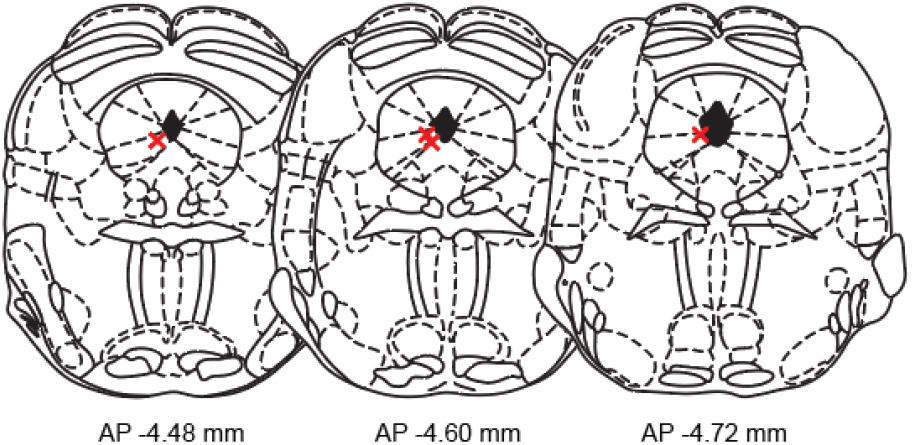
Schematic of the anatomical locations of GRIN lens implants in the 4 mice used in the experiments shown in Figure 3.

**Figure 4 – figure supplement 1.**
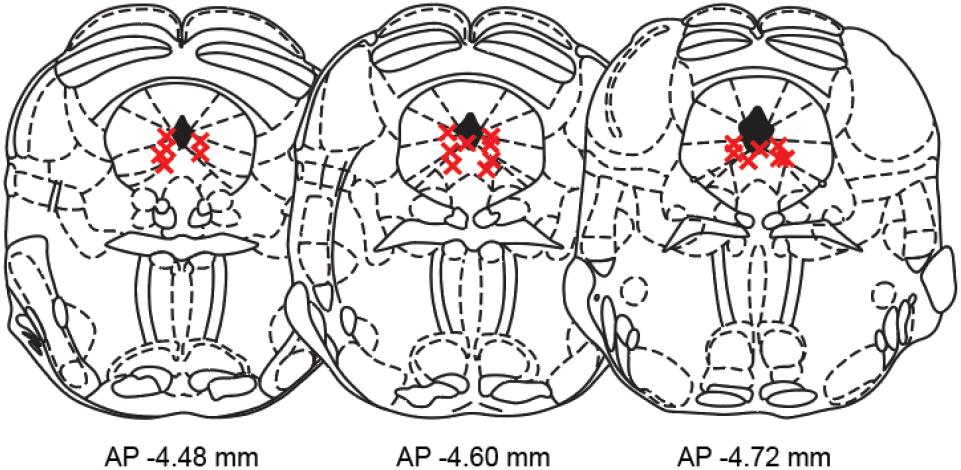
Schematic of the anatomical locations of the optical fiber implants in the 21 mice used in the experiments shown in Figure 4.

**Figure 4 – figure supplement 2.**
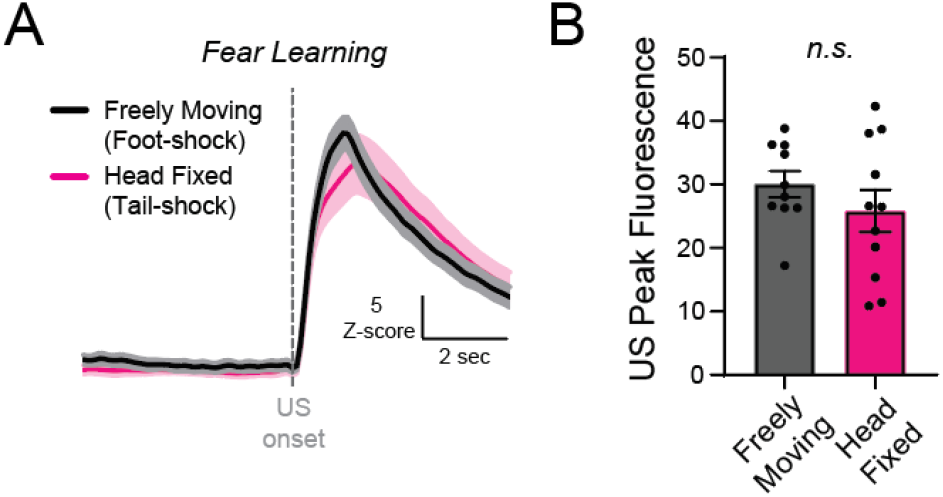
**(A)** Photometry responses to foot-shocks (in freely-moving mice, black) and tail-shocks (in head-fixed mice, magenta), averaged across all trials and mice. **(B)** Responses to foot-shocks (freely-moving mice, black) and tail-shocks (head-fixed mice, magenta) were not significantly different (n = 10 freely-moving mice, n = 11 head-fixed mice; unpaired t-test; t_19_ = 1.056, p = 0.3041). Data are presented as the mean ± S.E.M.

